# Computational cell cycle analysis of single cell RNA-Seq data

**DOI:** 10.1101/2020.11.21.392613

**Authors:** Marmar Moussa, Ion I. Măndoiu

## Abstract

The variation in gene expression profiles of cells captured in different phases of the cell cycle can interfere with cell type identification and functional analysis of single cell RNA-Seq (scRNA-Seq) data. In this paper, we introduce SC1CC (**SC1 C**ell **C**ycle analysis tool), a computational approach for clustering and ordering single cell transcriptional profiles according to their progression along cell cycle phases. We also introduce a new robust metric, Gene Smoothness Score (GSS) for assessing the cell cycle based order of the cells. SC1CC is available as part of the SC1 web-based scRNA-Seq analysis pipeline, publicly accessible at https://sc1.engr.uconn.edu/.

## 1 Background and motivation

The variation in gene expression profiles of single cells that are captured in different phases of the cell cycle can interfere with cell type identification and functional analysis of single cell transcriptomic data. In particular, it is important to differentiate between cell type and cell cycle effects when analyzing single cell RNA-Seq data. A first challenge in the computational analysis of cell cycle effects in single cell transcriptomics is to differentiate between cells that are actively proliferating and those that are quiescent, i.e., cells that do not actively divide but retain the ability to re-enter a proliferative state. A second computational challenge is to correctly label individual cells or cell clusters according to their phase in the cell cycle. The main cell cycle phases are G1 (where metabolic changes prepare the cell for division), S (where DNA synthesis replicates the genetic material), G2 (where molecular components needed for mitosis and cytokinesis are assembled), and M (where a nuclear division followed by cytokinesis occurs), although transition phases G1/S and G2/M are also commonly identified [6]. Such cell labels coupled with existing biological knowledge of genes associated with each of the cell cycle phases can assist functional analysis of single cell transcriptional profiles and interpretation of unsupervised scRNA-Seq clustering results. Finally, a third computational challenge is to order individual cells according to their progression along the cell cycle.

Although there are several existing methods for cell cycle analysis of single cell RNA-Seq data, most of them attempt to address one of the above-mentioned challenges in isolation. Our proposed SC1CC method enables a comprehensive analysis of the cell cycle effects that can be performed independently of cell type/functional annotation, hence avoiding hazardous manipulation of the single cell transcription data that could lead to misleading analysis results. Specifically, SC1CC can be used to distinguish proliferating from quiescent cells and provides the ability to annotate cell populations based on the cell cycle phase. Additionally, the cells are also ordered based on their progression along the cell cycle phases.

In the remainder of this section we briefly review some representative methods for individually addressing the above challenges in cell cycle analysis of scRNA-Seq. In Section 2 we introduce the datasets used in evaluation experiments and detail the computational methods underpinning SC1CC. In Section 3 we present and discuss experimental results comparing SC1CC with previous methods on real scRNA-Seq datasets of varying size and complexity and with experimental ground truth determined at different levels of resolution. Finally, we conclude in Section 4 with directions for future research.

### 1.1 ccRemover

The ccRemover tool [3] attempts to *remove* the cell-cycle effects from the single cell transcriptional profiles. This is done by identifying those principal components that, based on their loadings, capture mostly cell cycle effects in a low dimensional principal component analysis (PCA) projection of the scRNA-Seq data. Subtracting these components is expected to enhance gene expression variation due to differences in cell type. We performed an initial test to determine the effectiveness of ccRemover at removing cell cycle effects by running it with default settings on a dataset consisting of a 50%–50% mixture of Jurkat and 293T single cells that was previously profiled using the 10x Genomics droplet-based scRNA-Seq platform [1]. This dataset is comprised of cells of two different types (T lymphocyte and human embryonic kidney cells) that are well separated according to their original scRNA-Seq profiles (Supplementary Figures S1a) and b). However, after processing the scRNA-Seq data using ccRemover the two cell types appear nearly indistinguishable in the 3D t-SNE plot (Supplementary Figure S1c). This suggests that attempting to subtract the cell cycle signal using ccRemover without careful parameter tuning could result in inadvertently subtracting the cell type signal. For this reason, ccRemover was not included in further method comparisons in this paper.

### 1.2 scLVM and Cyclone

One of the earliest methods that address the cell cycle effect is the single-cell latent variable model (scLVM) algorithm [4] that uses a Bayesian latent variable model to reconstruct hidden factors in the expression profile of the cell-cycle genes. The scLVM algorithm assumes the first *k* factors to contain the cell-cycle effect and removes them from the dataset’s latent space. The assumption made by scLVM that all main signals provided by the cell cycle genes are exclusively cell cycle related was updated in a more recent method called Cyclone from the same authors [18]. Cyclone uses a classification algorithm based on selecting pairs of genes whose relative expression has a sign that changes with the cell-cycle phase in the training data. The learned gene pairs are used to quantify the evidence that a given cell is in one of three cell cycle phases (G1, S, or G2M). Specifically, under the recommended approach, Cyclone calculates for each cell a score between 0 and 1 for two of these phases, G1 and G2M. Cells with G1 or G2M scores above 0.5 are assigned to the G1 or G2M phases, respectively (if both scores are above 0.5, then the higher score is used to make the assignment). Cells with both G1 and G2M scores below 0.5 are assigned by default to the S phase. The method allows users to override these thresholds, but we used the recommended thresholds in our experiments. In Section 3 we present results comparing the accuracy of cell cycle labels inferred by Cyclone to those generated by SC1CC using datasets with both known and unknown cell cycle phase labels.

### 1.3 reCAT

The reCAT method [13] takes a different approach to cell cycle analysis. Rather than labeling the cells with an inferred cell cycle phase, reCAT attempts to *order* the cells in a manner consistent to their position along the cell cycle. The cell ordering problem is computationally modeled as a traveling salesman problem (TSP). First, reCAT performs normalization of the data followed by clustering of the cells. It then orders the identified clusters by finding a traveling salesman cycle. It also computes for each cell two scores (a Bayes score and a mean score) that differentiate between the cell cycle phases. Finally, a hidden Markov model (HMM) and a Kalman smoother are used to estimate the underlying gene expression levels of the ordered single cells. The results of experiments comparing the order reconstructed by reCAT to the order identified by SC1CC are presented in Section 3.

## 2 Methods

### 2.1 Datasets

In addition to the Jurkat-239T dataset described in Section 1.1 we used six other datasets to further evaluate the performance of SC1CC and existing cell cycle analysis tools. These datasets were selected to span a broad range of cell cycle related modalities. For example, all cells in the Human Embryonic Stem Cells (hESC) dataset described in Section 2.1 are expected to be proliferating, whereas the Peripheral blood mononuclear cells (PBMC) dataset described in Section 2.1 is expected to consist solely of quiescent cells [21]. The immune cells from anti-CTLA-4 treated mice (*α*-CTLA-4) dataset described in Section 2.1 and the mouse Hematopoietic Stem Cells (mHSC) from Section 2.1 are both expected to contain a mix of quiescent and proliferating cells. Finally, two all-dividing datasets with lower cell count are presented in Section 2.1: QuartzSeq data generated from embryonic stem (ES) cells and a dataset consisting of mouse embryonic stem cells (mESC). Three of these datasets (hESC, ES, and mESC) have labeled cell cycle phase annotations, while for the *α*-CTLA-4 and mHSC datasets only the percentage of proliferating cells was established in the original publications.

Basic quality control (QC) was uniformly applied to each of these datasets, whereby cells expressing less than 500 genes as well as genes detected in less than 10 cells were filtered out. Pre-processed versions of all datasets are accessible as example datasets for the SC1 web-based scRNA-Seq analysis pipeline [15], publicly available at https://sc1.engr.uconn.edu/.

#### Human embryonic stem cells (hESC, cycling cells)

There are very few scRNA-Seq datasets where the cell-cycle phase of each cell is known *a priori*. For this work, we used a labeled dataset of undifferentiated H1 human embryonic stem cells (hESCs) from [12]. Fluorescent ubiquitination-based cell-cycle indicator H1 (H1-Fucci) human embryonic stem cells were sorted according to the G1, S, and G2/M cell cycle phases by fluorescence activated cell sorting (FACS). Full-length scRNA-Seq data was generated for a total of 247 H1-Fucci cells (91 G1, 80 S, and 76 G2/M cells, respectively) captured using the Fluidigm C1 microfluidic platform.

#### Peripheral blood mononuclear cells (PBMC, non-cycling cells)

The PBMC dataset is comprised of a mixture of mature FACS-sorted dendritic cells, natural killer, B and T cells from a healthy donor from [21] and further analyzed in [14]. This dataset consists of 2,882 cells randomly sampled from seven PBMC sub-populations independently sorted by FACS. scRNA-Seq data for these cells was generated using the 10x Genomics droplet-based platform and the 3’-end v1 protocol, as described in [21]. Figure 1a shows a 3-dimensional *t-Distributed Stochastic Neighbor Embedding (t-SNE)* plot of the PBMC dataset and the breakdown into the seven cell types. Since PBMCs typically differentiate in the thymus or lymph nodes, this dataset is expected to contain only non-cycling cells.

**Fig. 1.**
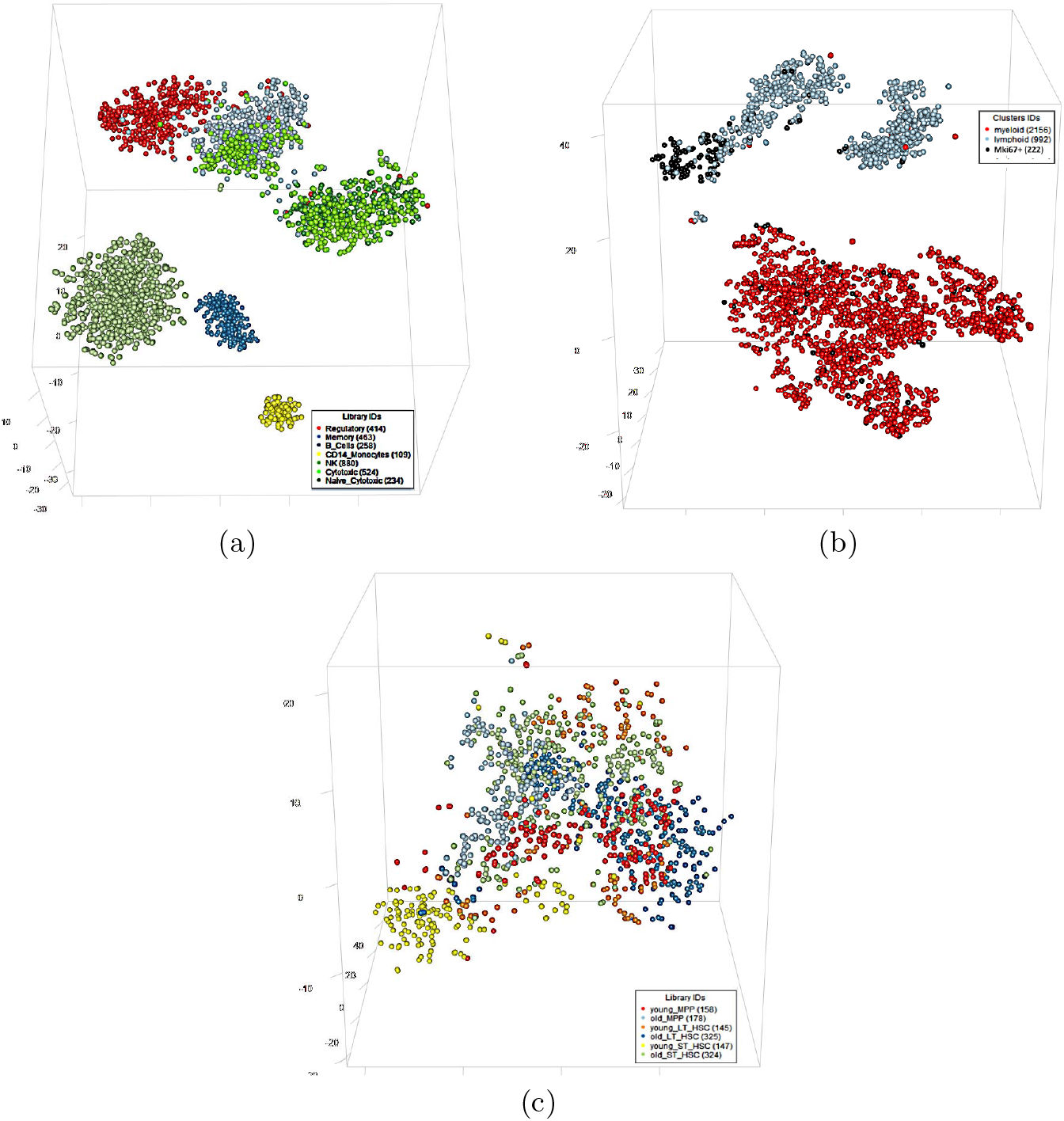
3-dimensional t-SNE plots of the PBMC, *α*-CTLA-4 and mHSC datasets. (a) 3D t-SNE plot of the 10x Genomics PBMC dataset consisting of 2,882 cells randomly sampled from seven PBMC sub-populations independently sorted by FACS. (b) 3D t-SNE plot of the α-CTLA-4 dataset consisting of 992 lymphoid (blue) and 2156 myeloid cells (red). The ‘Mki67-Hi’ cells (black) are a mixture of proliferating CD4+ T cells, CD8+ T cells, Tregs, and NK cells (≈ 17.5% of the lymphoid cells. (c) 3D t-SNE plot of the mHSC dataset consisting of a total of 1,277 MPP, ST-HSC, and LT-HSC cells, further grouped by the age of the mice (young and old).

#### Tumor infiltrating immune cells from anti-CTLA-4 treated mice (*α*-CTLA-4, mixture of cycling and non-cycling cells)

This dataset (publicly available in the NCBI GEO database under accession GSM3371686) was also generated using the 3’-end v1 scRNA-Seq protocol on the 10x Genomics platform. CD45+ cells were sorted by FACS from cell suspensions of dissociated tumors excised from mice treated with 9D9, an anti-CTLA-4 antibody, as described in [8]. According to the analysis in [8], this dataset, henceforth referred to as *α*-CTLA-4, consists of 992 lymphoid and 2,156 myeloid cells. Notably, the unsupervised clustering analysis of the *α*-CTLA-4 dataset in [8] has identified a cluster, labeled ‘Mki67-Hi’, comprised of a mixture of proliferating CD4+ T cells, CD8+ T cells, Tregs, and NK cells (≈ 17.5% of the lymphoid cells, see Figure 1b). Thus, this dataset is well suited for assessing the ability of our method to correctly differentiate between quiescent and proliferating cells.

#### Short- and long-term mouse hematopoietic stem cells from young and old mice (mHSC, mixture of cycling and non-cycling cells)

This scRNA-seq dataset (1,277 cells after applying QC) is publicly available in the NCBI GEO database under accession GSE59114. The dataset was used in [10] to dissect the variability in hematopoietic stem cell (HSC) and hematopoietic progenitor cell populations from young and old mice. A 3D t-SNE projection of the mHSC dataset is shown in Figure 1c. Based on the analysis in [10], this dataset is comprised of cells of three different types – Multipotent Progenitor Cells (MPP), short-term hematopoietic stem cell (ST-HSC), and long-term hematopoietic stem cell (LT-HSC) – that are further grouped by the age of the mice (young and old). The six cell populations are thoroughly analyzed in [10] with regards to cell cycle effect on differentiation while aging. We use the findings of this analysis as the ground truth for evaluating the performance of our approach. Specifically, the computational and biological analysis in [10] identifies 65% of all cells analyzed as non-dividing and estimates an equal percentage of proliferating cells in young and old mice for MPP and ST-HSC cells but not for LT-HSCs (of which old mice have fewer dividing cells). The analysis in [10] also estimates the percentages of cells in G1, S and G2M phases as 20%, 6% and 9% of the total, respectively.

#### Other datasets

To further test the robustness of our method, we used two other labeled datasets: QuartzSeq data generated from embryonic stem cells and analyzed in [17] (dataset ES) and mouse embryonic stem cells from [4] (dataset mESC). The QuartzSeq ES dataset is comprised of 8 cells labeled G1, 8 cells labeled M, and 7 cells labeled S phase, for a total of 23 cells. The mESC data is a filtered normalized FPKM dataset of 182 cells used as a training set for the Cyclone classifer in [4]. The results for these relatively small datasets are given in Supplementary Materials section.

### 2.2 The SC1 cell cycle (SC1CC) analysis tool

A repeated observation in single cell RNA-Seq data analysis is that a bias can be introduced by cell cycle effects. Indeed, such effects result in significant factor loadings of annotated cell cycle genes to the first few principal components for many scRNA-Seq datasets. Furthermore, it has been shown that the first few principal components obtained by using expression levels of annotated cell cycle genes are sufficient for capturing cell to cell similarities and the covariance due to cell cycle effects [4, 3, 18, 12, 7]. We leverage this observation in SC1CC and start by computing the first few principal components (PCs) for the sub-matrix of normalized scRNA-Seq counts comprised of cell cycle genes only.

The SC1CC implementation available at https://sc1.engr.uconn.edu/ allows users to select one of three different gene lists: the genes annotated with the “cell cycle” term (GO:0007049) in the Gene Ontology database [5], genes included in the Cyclebase 3.0 database of cell cycle related genes [16], and finally the list of periodic genes identified from single cell data in [7]. All results in this paper are based on the GO-annotated cell cycle gene unless otherwise indicated. The selected list of cell cycle genes is further filtered based on the gene expression values in the current dataset in order to keep only those expressed genes that have a correlation higher than *α* to at least one other cell cycle gene. The purpose of this step is to remove genes that – although annotated as cell cycle genes – do not have expression levels correlated with that of other cell cycle genes, and hence might represent outliers. Experimental results presented in Supplementary Table S1 suggest that cell cycle phase classification accuracy is relatively insensitive to the choice of correlation threshold *α*. All experiments reported in Section 3 were generated using the default value of 0.25 for *α*.

Since using a large number of PCs can add unnecessary noise to subsequent analysis steps, by default SC1CC automatically determines the number of relevant PCs by assessing the drop in variance explained for each pair of consecutive principal components. The online SC1CC implementation allows users to manually specify the number of PCs if desired. The principal component analysis is followed in SC1CC by a 3-dimensional t-SNE projection using the identified principal components. Performing t-SNE based dimensionality reduction using the main PCs aims to capture the local similarity of the cells without sacrificing the global variation already captured by the PC analysis. Next, the cells – now identified by their representation in t-SNE space – are clustered into a hierarchical structure (dendrogram) based on their Cosine similarity. Unless otherwise indicated all results reported in the paper are based on using hierarchical clustering with average linkage; the online SC1CC implementation also allows users to select between average linkage and Ward’s method.

Since the cell cycle is typically divided into 6 distinct phases (G1, G1/S, S, G2, G2/M, and M, see, e.g., [6]), by default SC1CC attempts to extract up to 7 clusters from the hierarchical clustering dendogram – corresponding to the 6 cell cycle phases plus at least one potential cluster of non-cycling cells – with a minimum cluster size threshold of 25 cells. The maximum number of clusters can be modified by the user in the online implementation of SC1CC, which also includes an ‘auto’ option for determining the optimal number of clusters based on the Gap Statistics Analysis algorithm from [19].

Finally, to generate an order of cells consistent to their position along the cell cycle, SC1CC reorders the leaves of the hierarchical clustering dendogram (corresponding to the individual cells) by using the *Optimal Leaf Ordering* (*OLO*) algorithm [2] as implemented in [9]. Performing additional leaf-node reordering is equivalent to minimizing the length of a Hamiltonian path [2]. For *n* cells, the dendrogram produced by the hierarchical clustering algorithm (essentially a rooted binary tree) has *n* – 1 internal nodes and 2^*n*−1^ possible leaf orderings. That is, at each internal node the left and right subtrees can be independently flipped or not. The OLO algorithm produces a leaf ordering that minimizes the sum of distances between adjacent leaves. The time complexity of the implementation in [9] is *O*(*n*^3^), and its practical performance as part of SC1CC is further improved since the pairwise distances between cells are already available from the distance based hierarchical clustering step.

#### Cluster Mean-Scores

Six groups of genes (G1, G1/S, S, G2, G2/M, and M genes, respectively) are formed by including cell cycle genes that are known to reach their peak expression in the corresponding cell cycle phases [13]. For each of these genes and each cell, a ‘z-score’ is computed by subtracting the gene’s mean expression level from the expression level of the gene in the cell and then dividing by the gene’s standard deviation. For each group of genes and each cluster identified during the hierarchical clustering step we compute a mean-score by averaging over cells in the cluster and genes in the group. The maximum mean-score of a cluster is used to determine its cell cycle phase. Note that with this procedure multiple clusters can be labeled with the same cell cycle phase, and some cell cycle phases may not be assigned as labels to any of the clusters. Also, since a mean-score of each gene group corresponding to each of the cell cycle phases can be calculated for each identified cluster, the maximum mean-score is relative between gene groups of different cell cycle phases and can only indicate a potential cell cycle phase designation. We therefore introduce in next sub-section an independent metric that can be used to distinguish dividing from non-dividing cells.

#### Gene-Smoothness Score (GSS)

Normalized gene scores computed as above or as defined by reCAT [13] or Cyclone [18] are relative between cell cycle phases and cannot distinguish clearly, if at all, between cycling vs. non-cycling cells or provide a useful metric for assessing cell orderings. We therefore propose a novel metric, referred to as Gene-Smoothness Score (GSS), based on serial correlation, i.e., the correlation between a given variable and a lagged version of itself. The GSS can be computed for any ordered group of cells and can help to directly assess the suggested cell order. Strengths of this metric include the fact that the cells do not need to have known cell-cycle labels and that no specific model assumptions are required for the marker gene expression (whether binary, bi-modal, sinusoidal, etc.). Our experiments also indicate that the GSS results are relatively insensitive to the choice of annotated cell cycle genes, hence the GSS can be useful even when a “perfect” annotation is not available.

The GSS of an ordered cluster/group *c* of cells is defined as

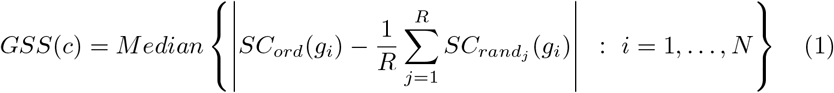

where *N* is the number of annotated cell cycle genes, *SC_ord_*(*gi*) denotes the first-order serial correlation of gene *i* with respect to the given cell order, and *SC_rand_j__*(*g_i_*), *j* = 1,…, *R*, denote the first-order serial correlation of gene i with respect to *R* randomized cell orders (we use R = 50 in all experiments). The first-order serial (or auto-) correlation is the correlation value between a given gene expression vector and a version of itself shifted by one position. Serial correlation is a value between −1 and 1. First-order serial correlation near 0 implies that there is no overall correlation between adjacent data points. On the other hand, a first-order serial correlation near 1 suggests a smoothly varying series, while a first-order serial correlations near −1 indicates a series that alternates between high and low values. Because individual cell cycle genes can be expressed in different patterns throughout the cell cycle phase transitions, and even abruptly switch direction when the assessed cluster includes mostly cells in one of the transient cell cycle phases (G1/S or G2/M), we define GSS as the median (over all cell cycle genes) of the absolute differences between the serial correlation of a gene’s expression values ordered according to the given cell ordering and the average serial correlation computed over *R* randomized orders.

A cluster/group of cells is considered to be cycling/dividing when its GSS is greater than an error margin *ε* (default 0.05), i.e., when at least 50% of the genes have an absolute difference in serial correlations between randomized order and identified cell cycle order of at least 0.05. The value of the error margin is set to 0.05 by default but can be adjusted by the user in the online SC1CC implementation. The GSS score is more robust with a higher number of cells per cell cycle cluster, as the chance of a random order producing spurious auto-correlation and therefore high GSS scores is lower when more data points are included in the series.

Figure 2 provides examples of cell cycle genes that contribute positive values to the GSS score in the hESC dataset and illustrates their expression values for both SC1CC and randomly ordered cells. The online implementation of SC1CC allows the user to select any cell cycle gene of interest and examine its normalized expression levels along the inferred order. In Figure 2, gray dots represent normalized gene expression values for individual cells, while the red and blue curves represent the fitted local polynomial regression of these values for the SC1CC and a random cell order, respectively. As expected, the fitted expression lines under random ordering of the cells convey no recognizable pattern and stay nearly flat close to an altitude of 0. In contrast, the SC1CC cell order results in fitted curves that appear to peak at different positions, consistent with these gene’s involvement in different cell cycle phases.

**Fig. 2.**
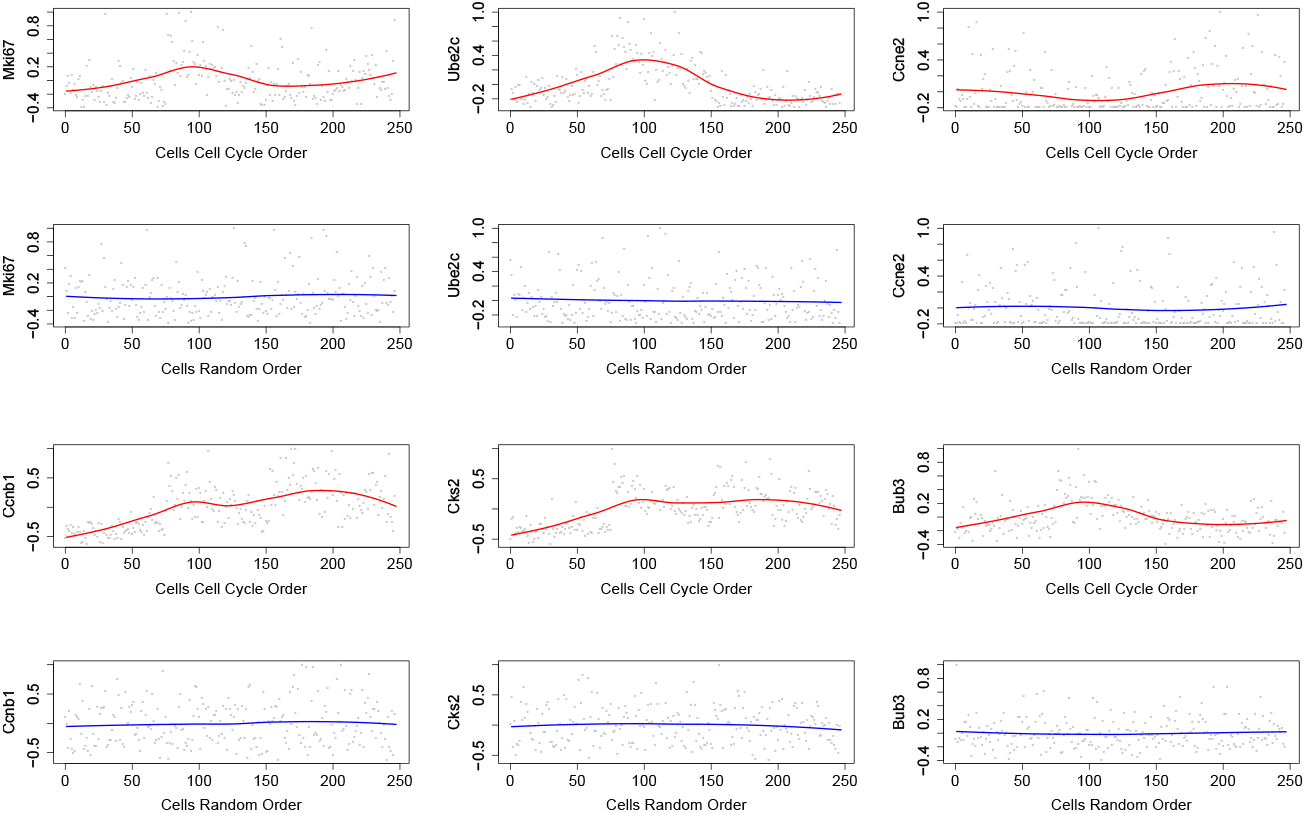
Example cell cycle genes in the hESC dataset. Normalized expression levels for select cell cycle genes and cells ordered by SC1CC (red) vs. a shuffled cells order (blue). Different cell cycle genes follow different patterns of expression along the cell cycle phases. Given the SC1CC inferred cell order, which reflects the cells’ progression through the cell cycle, different patterns for individual cell cycle genes can be seen for different genes associated with the cell cycle, including Mki67, Ube2c, Ccne2, Ccnb1, Cks2 and Bub3.

## 3 Results and Discussion

### 3.1 Results on the hESC Dataset

The cell order inferred by SC1CC’s OLO algorithm and the cell cycle order reconstructed by reCAT are shown in Figure 3a. SC1CC groups together almost all cells labeled with the same phase. Although the reCAT order maintains the grouping of G1 cells, cells from S and M phases are highly interleaved in this order. The SC1CC order also has a higher GSS score of 0.0632 compared to 0.0519 for the reCAT order.

**Fig. 3.**
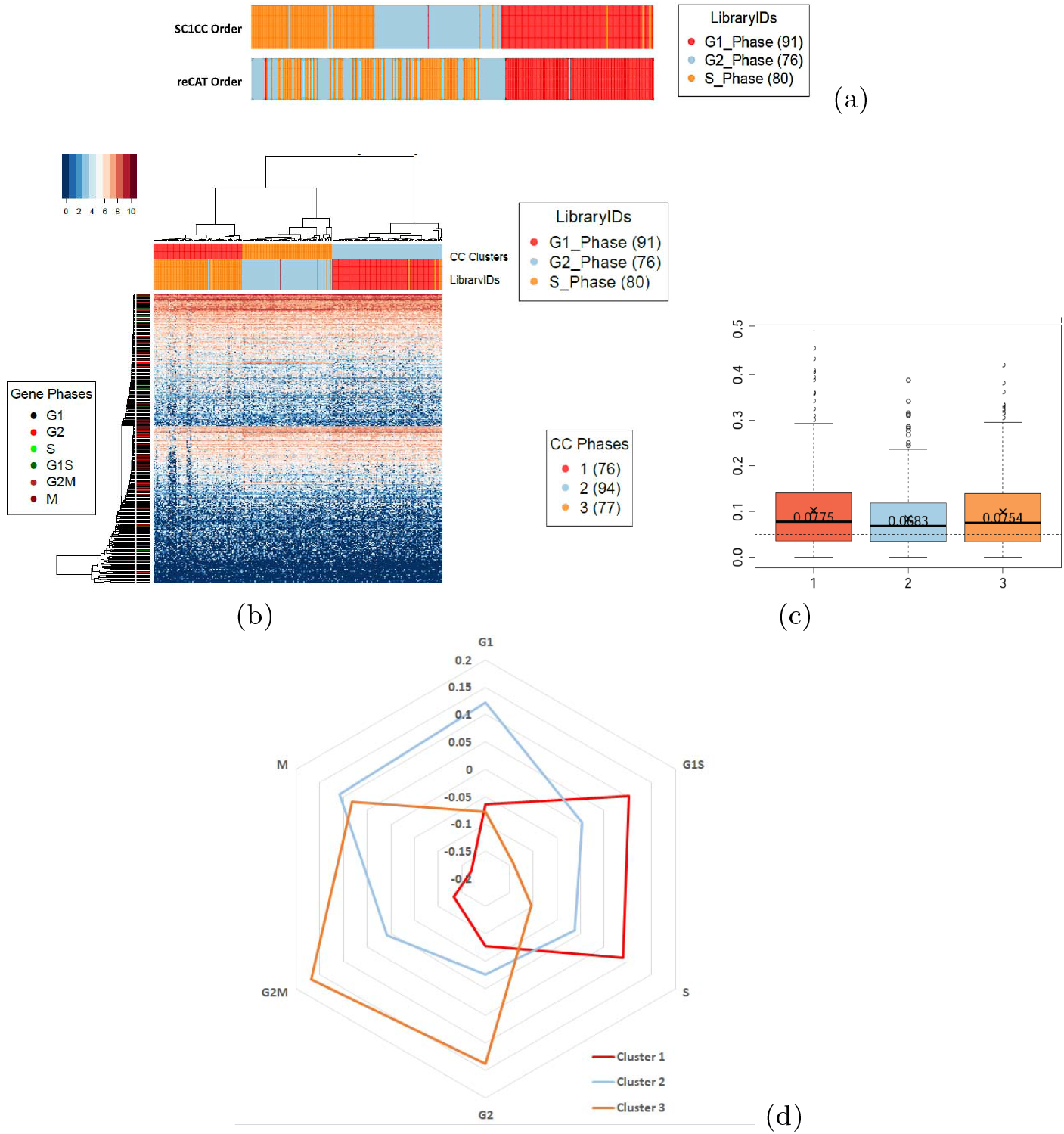
Cell cycle analysis of the hESC dataset. (a) hESC cells orders inferred by SC1CC and reCAT. Experimentally determined cell labels are color coded as library IDs. SC1CC groups the majority of cells from each phase together (G1 in red, G2 in blue, and S in orange), whereas only G1 cells are grouped together in the reCAT order. (b) Heatmap of *log2 (x* + 1) expression values of cell cycle genes for the hESC cells using ‘GO:00049’ gene list ordered according to SC1CC. Colors in the top bar labeled ‘CC Clusters’ represent the identified cell cycle clusters according to SC1CC, whereas the colors in the ‘Library IDs’ bar of the heat map indicate the cell cycle phases determined by FACS. (c) GSS for the three clusters identified by running SC1CC on the hESC cells. (d) Mean-scores for each of the three clusters identified by SC1CC and each of the six considered cell cycle phases. Based on majority matching of cell labels determined by FACS, the three clusters (Cluster 1 in red, Cluster 2 in blue and Cluster 3 in orange) are comprised of cells in the S, G1, and G2 phases respectively.

Figure 3b displays the heat map of *log_2_(x* + 1) expression values of cell cycle genes for the hESC cells ordered according to SC1CC. Colors in the top bar labeled ‘CC Clusters’ represent the identified cell cycle clusters according to SC1CC, whereas the colors in the ‘Library IDs’ bar of the heat map indicate in this case the cell cycle phases determined by FACS. Note that the colors for library IDs and inferred cell cycle clusters are assigned independently from the same color palette in our online implementation and therefore are not necessarily in one-to-one correspondence. The heat map in Figure 3b was generated by running SC1CC using the set of genes associated with the GO term “Cell Cycle” (GO:0007049); heat maps for Cyclebase 3.0 and periodic genes from [7] are given in Supplementary Figure S2. The hierarchical clustering algorithm implemented by SC1CC identifies three clusters. The GSS scores (Figure 3c) for the three clusters were 0.0775, 0.0683, and 0.0754, respectively, indicating that all clusters consist of dividing cells, as expected.

Figure 3d gives the mean scores for each of the three clusters identified by SC1CC and each of the six considered cell cycle phases. Based on majority matching of cell labels determined by FACS, the three clusters are comprised of cells in the S, G1, and G2 phases, respectively. Albeit not perfect, the cluster assignments based on peak mean scores have good agreement. Specifically, cluster 1 (consisting of S phase cells according to the FACS labels) has very close highest mean scores for the G1S and S phases, with the G1S score slightly higher. Cluster 2 (G1 according to FACS) has two close highest scores for G1 and M phases, with the G1 score slightly higher. Finally, cluster 3 (G2 according to FACS) has two close highest mean scores for the G2 and G2M phases, with the G2M score slightly higher. The relatively low number of cells as well as the limited resolution of the ground truth labels are both likely contributing factors to the near-ties in peak score assignments for the three clusters.

In Table 1 we compare the clusters (cell labels) generated by Cyclone with the clusters inferred by SC1CC using different cell cycle gene sets for the hESC dataset. We assess clustering accuracy using the macro and micro-accuracy measures from [11] and [20], defined as:

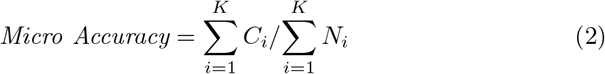

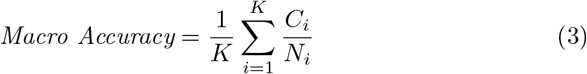

where *K* is the number of classes, *N_i_* is the size of class *i*, and *C_i_* is the number of correctly labeled samples in class *i* relative to the ground truth.

**Table 1.** Clustering accuracy for Cyclone and SC1CC run with three different gene lists on the hESC dataset.

|  | Cyclone | SC1CC |  |  |
| --- | --- | --- | --- | --- |
|  |  | Cyclebase 3.0 | GO | Periodic Genes |
| G1-Phase | **1.000** | 0.9890 | 0.9890 | **1.000** |
| G2-Phase | 0.9342 | 0.7895 | **0.9605** | 0.8290 |
| S-Phase | **1.000** | **1.000** | 0.9125 | 0.9875 |
| Micro Accuracy | **0.9798** | 0.9312 | 0.9555 | 0.9433 |
| Macro Accuracy | **0.9781** | 0.9262 | 0.9540 | 0.9388 |

Both Cyclone and SC1CC cluster the cells with high accuracy, with Cyclone scoring slightly higher. As detailed in Section 2.2, SC1CC gives the user the choice to use three different lists: genes included in the Cyclebase 3.0 database [16], genes annotated with the “cell cycle” term (GO:0007049) in the Gene Ontology database [5], and the list of periodic genes identified from single cell data in [7]. As can be seen in Table 1, the genes associated to the term “Cell Cycle” (GO:0007049) from the The Gene Ontology (GO) database achieve slightly higher clustering micro- and macro-accuracy for the hESC dataset compared to the other two gene sets.

### 3.2 Results on the PBMC dataset

As described in Section 2.1, the PBMC dataset is expected to include mostly non-dividing cells, which is confirmed by the results of the SC1CC analysis. Figure 4a shows the heat map of the PBMC cells featuring the GO cell cycle related genes that are expressed in the dataset (using *log*_2_ (*x* + 1) expression) and the clustering obtained by SC1CC. The majority of the genes have low expression levels in most cells. Furthermore, the GSS scores of all clusters fall below the 0.05 cutoff and hence they are all labeled as non-dividing by SC1CC (Figure 4b), as expected. Cyclone labels 2,192 of the 2,882 cells in the PBMC dataset as G1, 398 as G2M, and 292 as S phase cells, underscoring the need for a separate analysis step to determine if the cells are actually cycling.

**Fig. 4.**
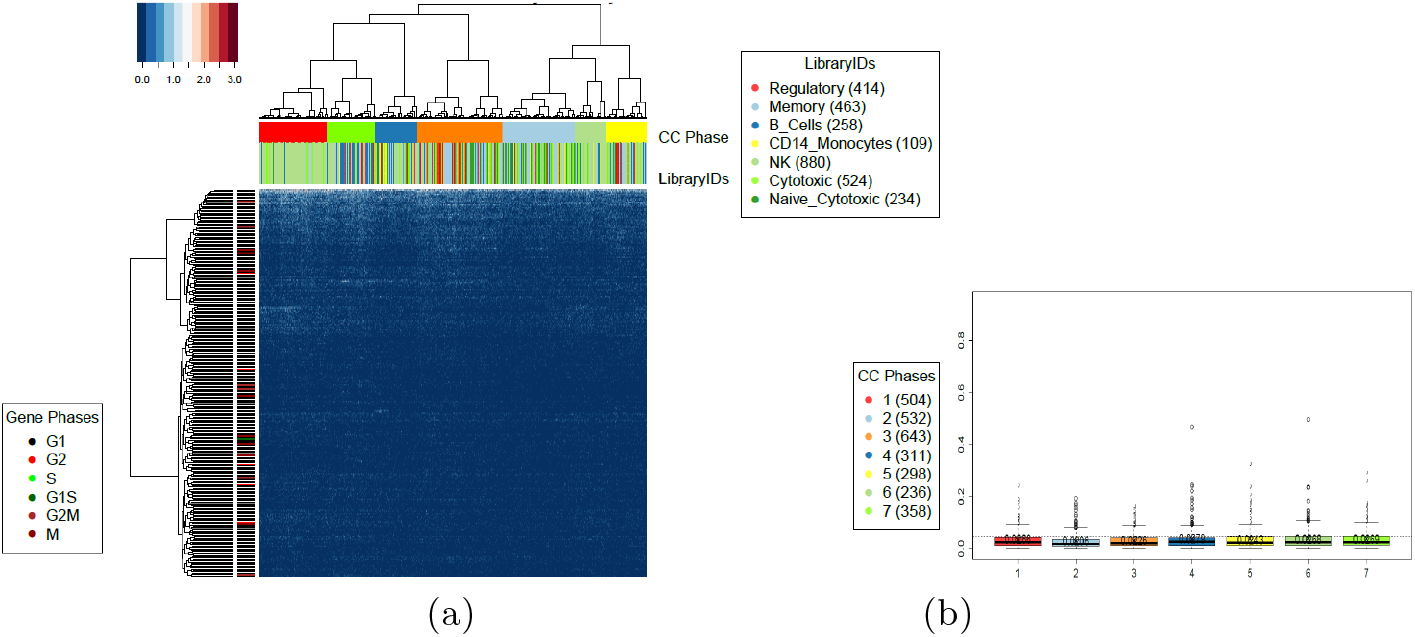
Cell cycle analysis of the PBMC dataset. Heat map and cell order (a) along with GSS scores for the clusters inferred by SC1CC (b) on the PBMC dataset. GSS scores for all clusters fall below a cutoff of 0.05 and are labeled as non-dividing by SC1CC.

### 3.3 Results on the *α*-CTLA-4 dataset

As discussed in Section 2.1, the *α*-CTLA-4 dataset is expected to include a mix of dividing and non-dividing cells. This is the most likely scenario for many scRNA-Seq datasets where no knowledge of the cell cycle effect within the data is available *a priori*. We reasoned that the best analysis approach for such data is to perform a two stage analysis, where we first separate the dividing from the non-dividing cells, followed by a detailed cell cycle analysis of the potentially dividing cells identified in the first step. Indeed, after clustering and ordering the cells using SC1CC, we are able to distinguish the potentially dividing cells by their GSS score. Figure 5a shows the *log*_2_ (*x* + 1) expression heat map of the 3,148 *α*-CTLA-4 cells passing the default QC described in Section 2.1 based on the GO cell cycle genes and using the first 4 principal components. One cluster (cluster 7 in light green color) consists of 193 cells that show markedly higher expression levels for the cell cycle genes. Independent clustering analysis based on full gene expression profiles performed using the SC1 pipeline shows that cluster 7 is comprised mostly of lymphoid cells (light blue in the horizontal bar labeled “Clusters” in the heat map). This cluster has the highest GSS score, exceeding the SC1CC detection threshold for dividing cells, as shown in Figure 5b. Indeed, this cluster closely matches the “Mki67-Hi” cluster identified in [8] as consisting of highly proliferative lymphoid cells. Further SC1CC analysis of the 193 cells in this cluster based on the Cyclebase 3.0 gene list reveals three sub-clusters (Figure 5c), all of which are found to be actively dividing according to GSS scores (Figure 5d). Cell cycle phase assignments based on maximum mean-scores suggests that the three sub-clusters consist of cells in the M, S, and G1S phases, respectively (Figure 5e). For the sake of completeness we also tested Cyclone classification method on the of the 3,148 α-CTLA-4 cells, and 2,957 cells were labeled as G1, 149 were labeled as G2M, and 42 were labeled as S phase cells.

**Fig. 5.**
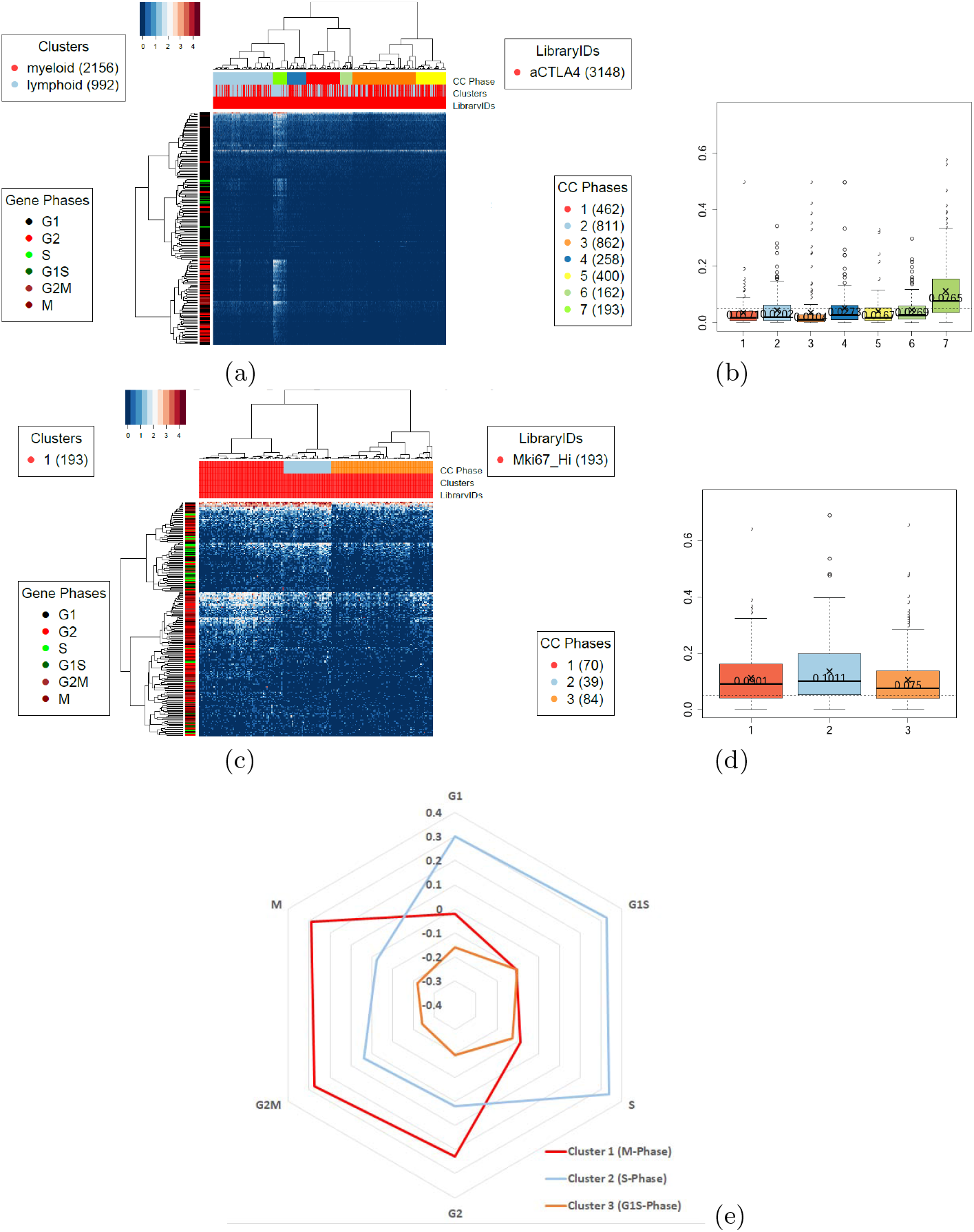
Cell cycle analysis of the *α*-CTLA-4 dataset. Heat map and cell order (a) along with GSS scores for the clusters inferred by SC1CC (b) on the *α*-CTLA-4 dataset. The 193 cells in cluster 7 are further partitioned by SC1CC into 3 sub-clusters (c), all of which are marked as actively dividing based on GSS scores (d). Mean-scores for each of the three sub-clusters of dividing cells and each of the six considered cell cycle phases are given in (e). Mean-scores for each of the three sub-clusters of dividing cells and each of the six considered cell cycle phases are given in (e). The maximum mean-scores of sub-clusters 1 (red), 2 (blue), and 3 (orange) are achieved for the M, S, and G1/S phases, respectively.

### 3.4 Results on the mHSC dataset

As discussed in Section 2.1, this dataset also includes a mix of dividing and nondividing cells. As with the *α*-CTLA-4 dataset analysis, we followed a two stage SC1CC analysis approach, where we first separate the dividing from the non-dividing cells, followed by a detailed cell cycle analysis of the potentially dividing cells identified in the first step. In excellent agreement with the percentages reported in [10], the first analysis stage (Figure 6a-b) places 472 of the 1,277 mHSC cells (36.96%) in a dividing cluster with GSS score of 0.3075, and the remaining cells in a non-dividing cluster with GSS score of 0.0285. Furthermore, as shown in Table 2, the percentage of dividing cells identified by SC1CC among the three cell types identified in [10] are indeed approximately equal in young and old mice, with the exception of long term HSC, only 13% of which are dividing in old mice compared to 35% in young mice.

**Fig. 6.**
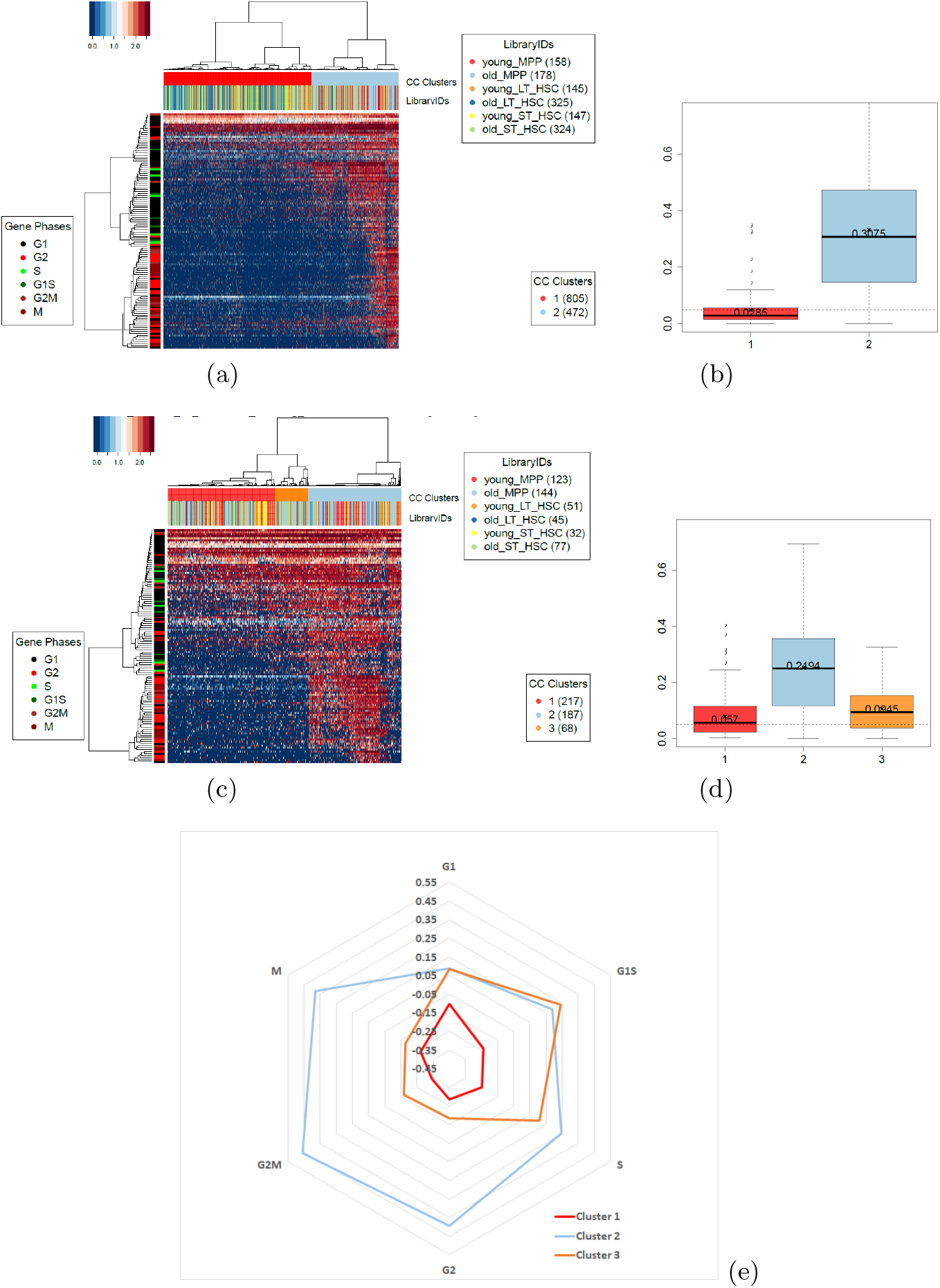
Cell cycle analysis of the mHSC dataset. Heat map and cell order (a) along with GSS scores for the clusters inferred by SC1CC (b) on the mHSC dataset. The 472 cells in cluster 2 are further partitioned by SC1CC into 3 sub-clusters (c), all of which are marked as actively dividing based on GSS scores (d). Mean-scores for each of the three sub-clusters of dividing cells and each of the six considered cell cycle phases are given in (e). The maximum mean-scores of sub-clusters 1 (red), 2 (blue), and 3 (orange) are achieved for the G1, G2M, and G1/S phases, respectively.

**Table 2.** The inferred numbers of dividing vs. non-dividing cells in the six cell populations of the mHSC dataset.

|  | Non-Dividing | Dividing | % Dividing |
| --- | --- | --- | --- |
| Old mice MPP | 34 | 144 | 80% |
| Young mice MPP | 35 | 123 | 78% |
| Old mice ST-HSC | 247 | 77 | 24% |
| Young mice ST-HSC | 115 | 32 | 22% |
| Old mice LT-HSC | 280 | 45 | 13% |
| Young mice LT-HSC | 94 | 51 | 35% |

The analysis in [10] ‘roughly’ estimates the percentage of cells in G1, G1/S and G2M phases as 20%, 6% and 9% respectively. Following the second stage of analysis, where cycling cells identified in first stage are further clustered and assigned cell cycle phases based on peak mean scores (Figure 6c-e), SC1CC identifies 217 cells (17% of total) as G1, 68 cells as G1/S (5.3% of total), and 187 cells as G2M (14.6% of total).

## 4 Conclusion

In this paper we introduce SC1CC (see Supplementary Figure S5 for a high level workflow), a novel method for clustering and ordering single cell transcriptional profiles according to their cell cycle phase. The main contributions include a novel technique for ordering cells based on hierarchical clustering and optimal leaf ordering, and a new GSS metric based on serial correlation for assessing gene expression change smoothness along a reconstructed cell order as well as differentiating between cycling and non-cycling groups of cells. While many of the existing methods focus on a specific aspect of scRNA-Seq cell cycle analysis (e.g., assigning phase labels, ordering the cells, or removing the cell cycle contribution to gene expression), SC1CC is, to our best knowledge, the first method that enables a comprehensive analysis of the cell cycle effects, addressing four complementary analysis aspects. SC1CC differentiates between dividing and non-dividing cells, clusters the cells based on cell cycle effects independently from cell type effects, while also assigning cell cycle phases to the resulting clusters and ordering the cells based on their progression along the cell cycle phases. SC1CC has been implemented in R and deployed via a user-friendly interactive interface as part of the SC1 scRNA-Seq analysis pipeline, freely accessible at https://sc1.engr.uconn.edu/.

Empirical evaluation experiments on a diverse set of real scRNA-Seq datasets show that the GSS robust evaluation metric which allows distinguishing with high accuracy between dividing and non-dividing cells based on minimal assumptions about the underlying cell cycle gene expression changes. In direct comparisons with the existing specialized tools, SC1CC also achieves similar or better accuracy for clustering the cells according to cell cycle phases or ordering them according to the progression along the cell cycle phases. Importantly, SC1CC analysis is performed orthogonally to cell type identification, avoiding potentially artifacts of sequential analysis advocated in [3].

Currently SC1CC relies on prior biological knowledge in the form of annotated cell cycle gene lists. A possible future research direction is to improve accuracy by augmenting prior annotations with cell cycle genes identified from the data itself based on their expression pattern along the cell-cycle ordering of the cells.

## Supplementary Material

### Results on QuartzSeq ES [17] and mESC[4] datasets

To further test the robustness of our method, we used the datasets from [17] (dataset QuartzSeq ES) and the mouse ESC from [4] (dataset mESC).

QuartzSeq ES data is a small dataset with 8 cells labeled G1, 8 cells labeled M and 7 cells labeled S phase, a total of 23 cells. Although it contains less cells than the recommended minimum size for SC1CC method SC1CC ordered the cells with perfect accuracy of 1 using the Cyclebase gene list and average linkage in the HC step. As this is a small set, only two clusters are reported by the online implementation of SC1CC. Results of the analysis are shown in Figure S3. To compare our results, and using Cyclone classifier method for this set (using Cyclone’s default settings), 6 were labeled G1, 8 in M and 10 in S phase, with overall accuracy of 0.87.

The mESC data is a filtered FPKM set of 182 cells normalized by [4] and used as the training set for Cyclone classifier, hence the accuracy for Cyclone method for this set was 0.93. Although it is unclear how the normalization from [4] affects SC1CC, the overall accuracy is 0.85 using 5 PCs, the Periodic gene list and Ward algorithm in the HC step of SC1CC, details of this analysis are shown in Figure S4.

## Acknowledgments

This work was partially supported by NSF Award 1564936, NIH grants 1R01MH112739-01 and 2R01NS073425-06A1, and a UConn Academic Vision Program Grant.

---

**Fig. S1.**
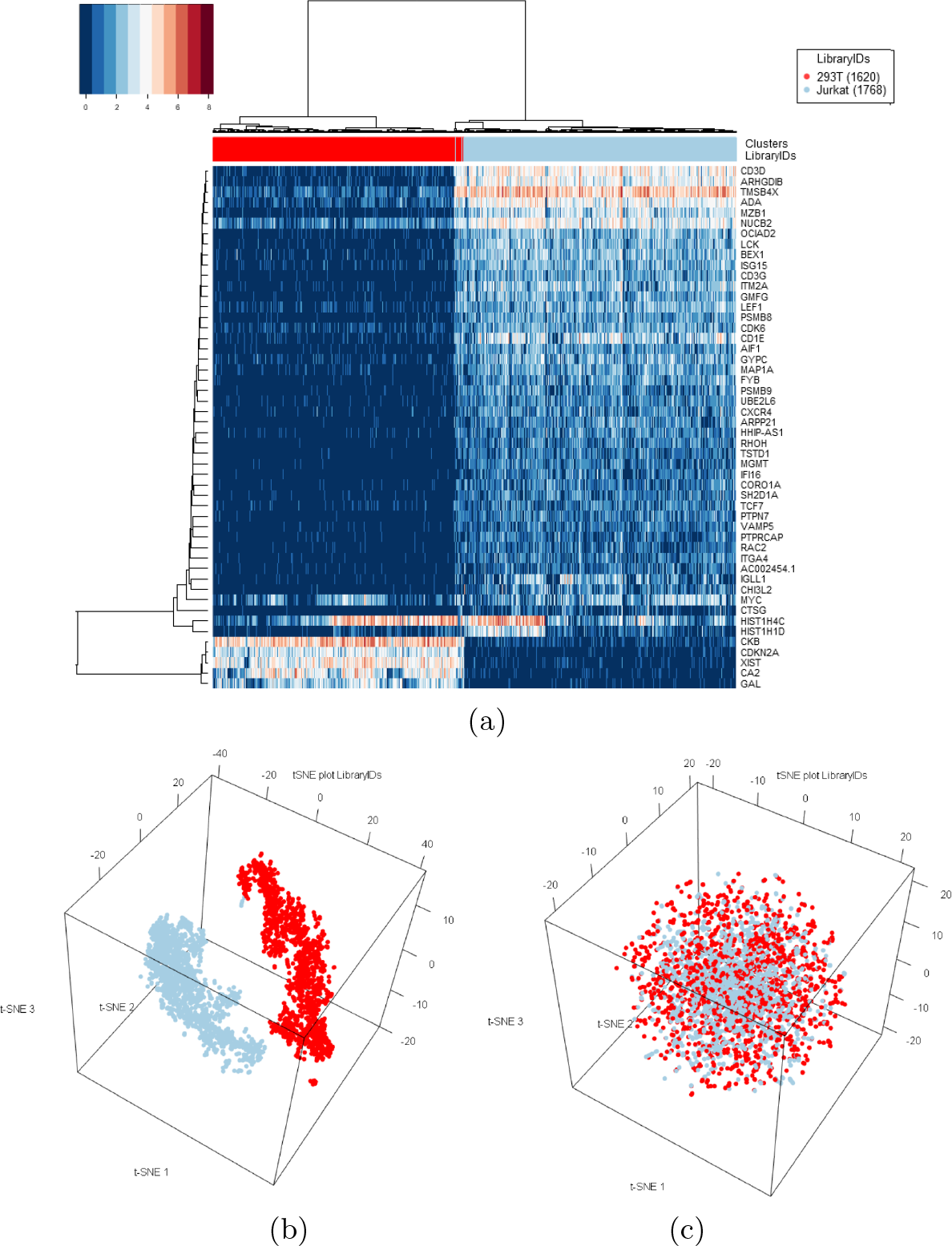
Jurkat-239T data. a) Heat map of Jurkat-239T data showing blue and red colors distinguishing the Jurkat and 239T cells respectively.We used TF-IDF based Hierarchical Clustering method from [14] and the heat map features the top 50 genes with highest average TF-IDF scores. Both markers, CD3D and XIST that are used in [21] to identify the cell lines in the mixture are selected amongst the list of genes with highest TF-IDF scores. **ccRemover effect on data variability.** b) 3D t-SNE plot of Jurkat-239T data (Blue and red colors distinguish the Jurkat and 239T cells respectively). c) 3D t-SNE plot of the Jurkat-293T dataset after applying ccRemover with default settings. As in [21] we inferred the cluster/library labels based on the expression of cell type-specific markers, The blue cluster corresponds to Jurkat cells (preferentially expressing CD3D), and red corresponds to 293T cells (preferentially expressing XIST, as 293T is a female cell line, while Jurkat is a male cell line).

**Fig. S2.**
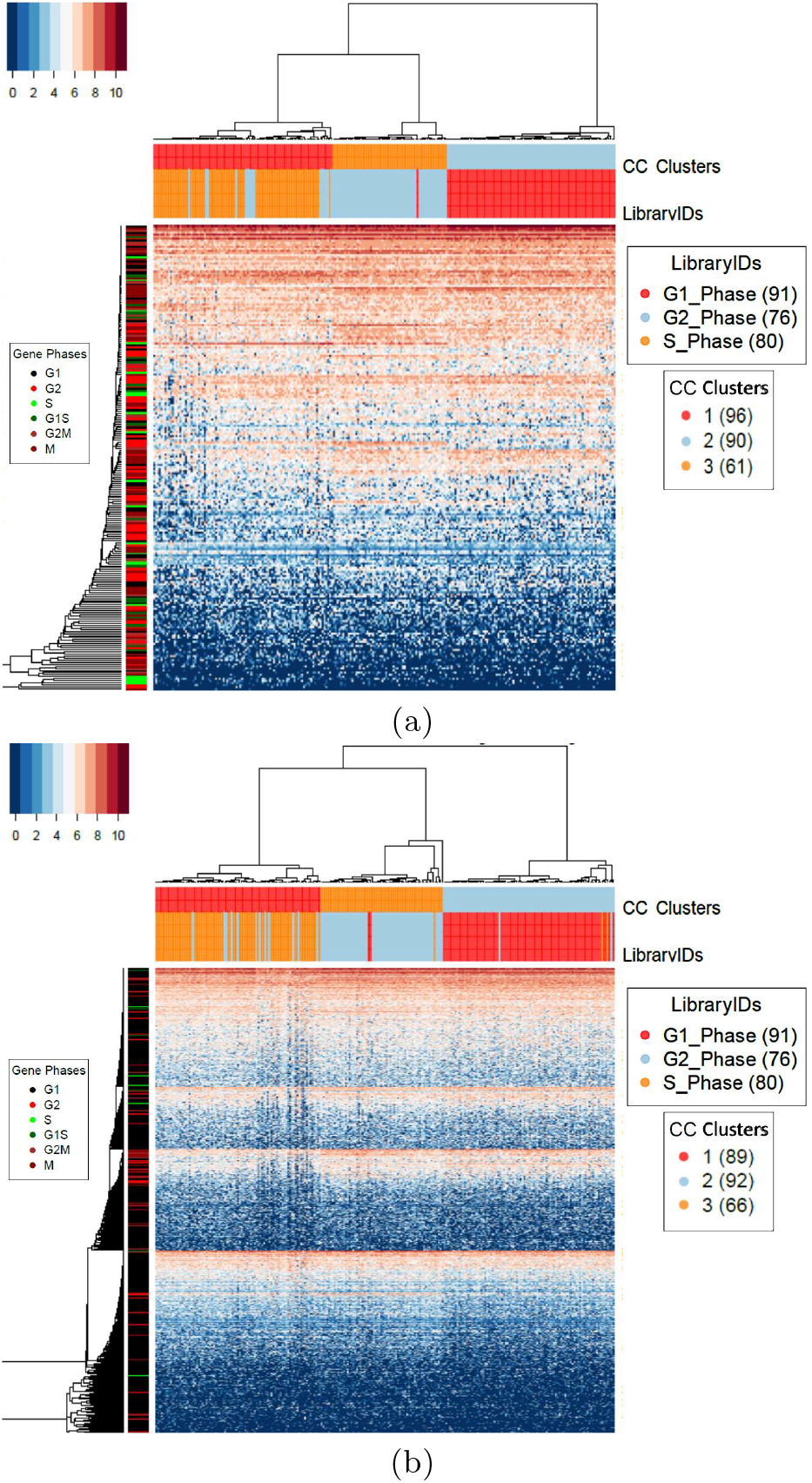
Different gene lists can affect the cell cycle analysis. a) Heat map of hESC database using Cyclebase 3.0 genes list. b) Heat map of hESC database using periodic gene list from [7].

**Table S1.** Correlation filter applied to hESC dataset

| Correlation filter | 0 | 0.05 | 0.1 | 0.15 | 0.2 | 0.25 | 0.3 | 0.35 |
| --- | --- | --- | --- | --- | --- | --- | --- | --- |
| Number of Genes | 580 | 580 | 580 | 580 | 569 | 429 | 250 | 146 |
| G1 | 0.930 | 0.930 | 0.930 | 0.930 | 0.930 | 0.989 | 0.950 | 0.960 |
| G2 | 0.900 | 0.900 | 0.900 | 0.900 | 0.900 | 0.961 | 0.800 | 0.800 |
| S | 0.980 | 0.980 | 0.980 | 0.980 | 0.980 | 0.913 | 0.980 | 0.940 |
| Macro Accuracy | 0.937 | 0.937 | 0.937 | 0.937 | 0.937 | 0.954 | 0.910 | 0.900 |
| Micro Accuracy | 0.927 | 0.927 | 0.927 | 0.927 | 0.927 | 0.956 | 0.919 | 0.907 |

**Fig. S3.**
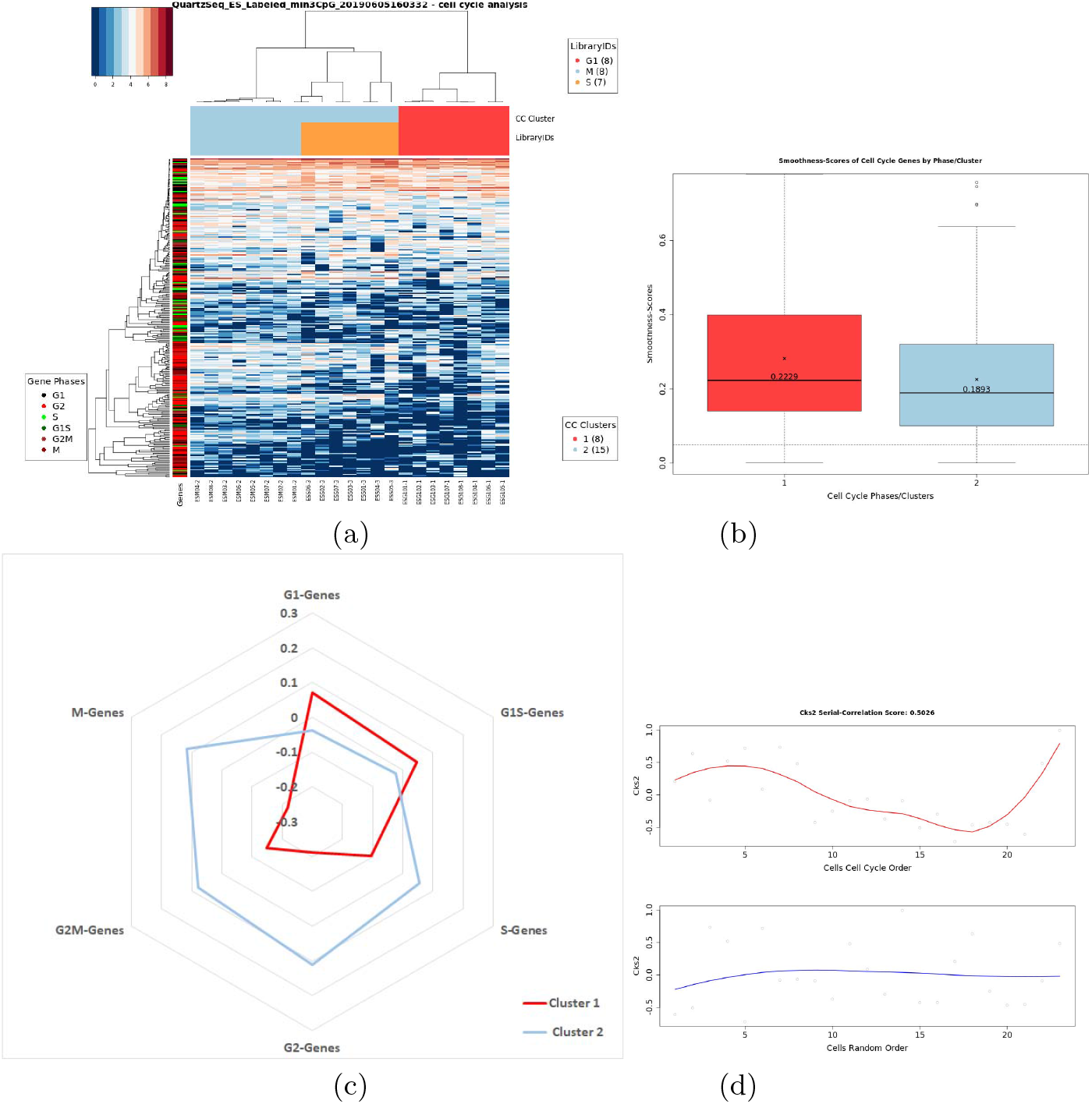
QuartzSeq ES data. Heat map and cell order (a) and GSS scores for the clusters inferred by SC1CC on the QuartzSeq ES dataset. The mean scores for each cell cycle cluster identified by SC1CC is given for each of the six considered cell cycle phases, the peak value of the mean-score of Cluster 1 (red) is found in G1 phase, Cluster 2 (blue) in G2M phase and Cluster 3 (orange) in G1/S phase (c) and as an example, Normalized expression levels for Ccnb2 of cells ordered by SC1CC (red) vs. a shuffled cells order (blue) in (d)

**Fig. S4.**
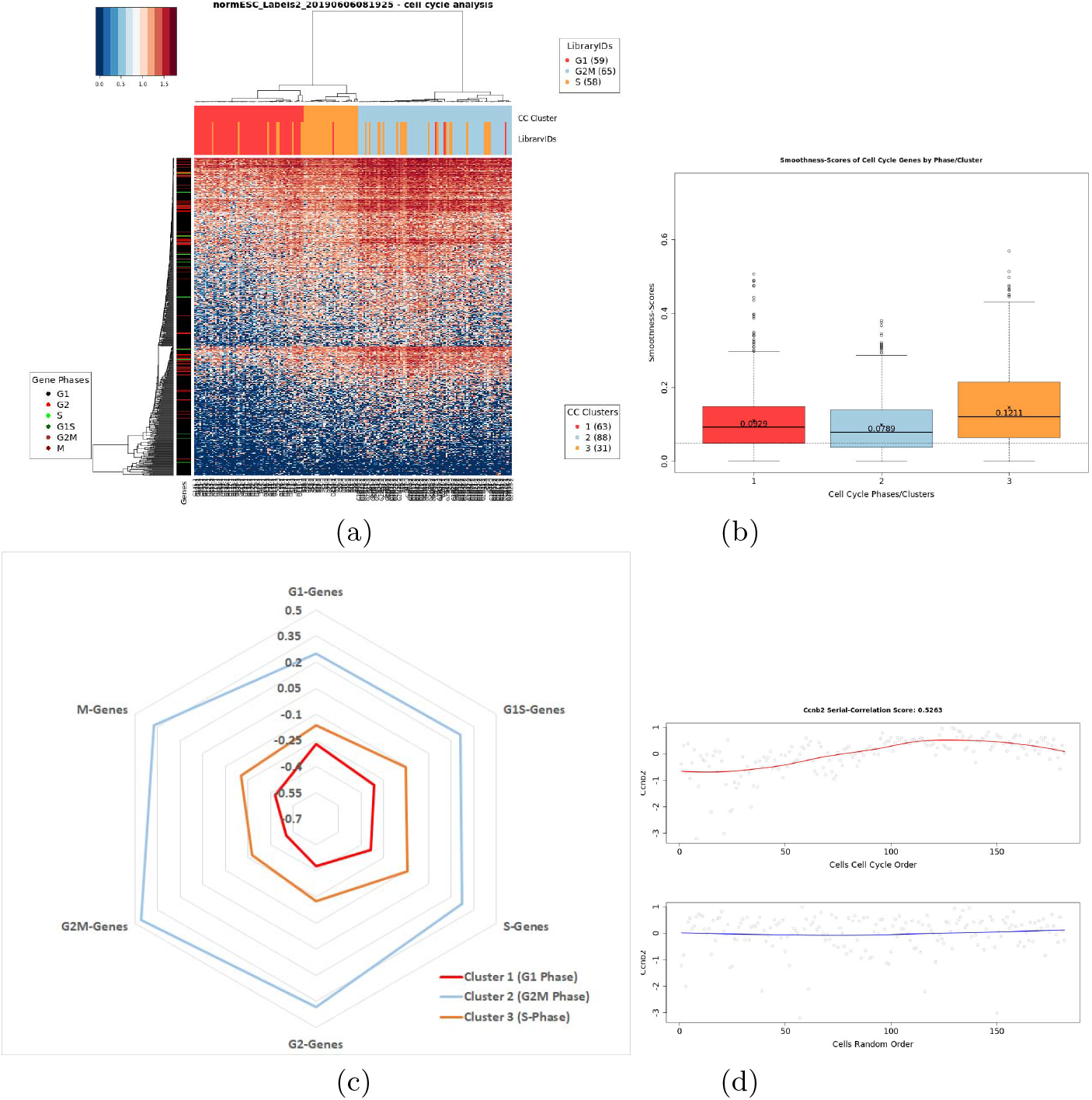
mES data. Heat map and cell order (a) and GSS scores for the clusters inferred by SC1CC on the mES dataset. The mean scores for each cell cycle cluster identified by SC1CC is given for each of the six considered cell cycle phases, the peak value of the mean-score of Cluster 1 (red) is found in G1 phase, Cluster 2 (blue) in G2M phase and Cluster 3 (orange) in G1/S phase (c) and as an example, Normalized expression levels for Ccnb2 of cells ordered by SC1CC (red) vs. a shuffled cells order (blue) in (d).

**Fig. S5.**
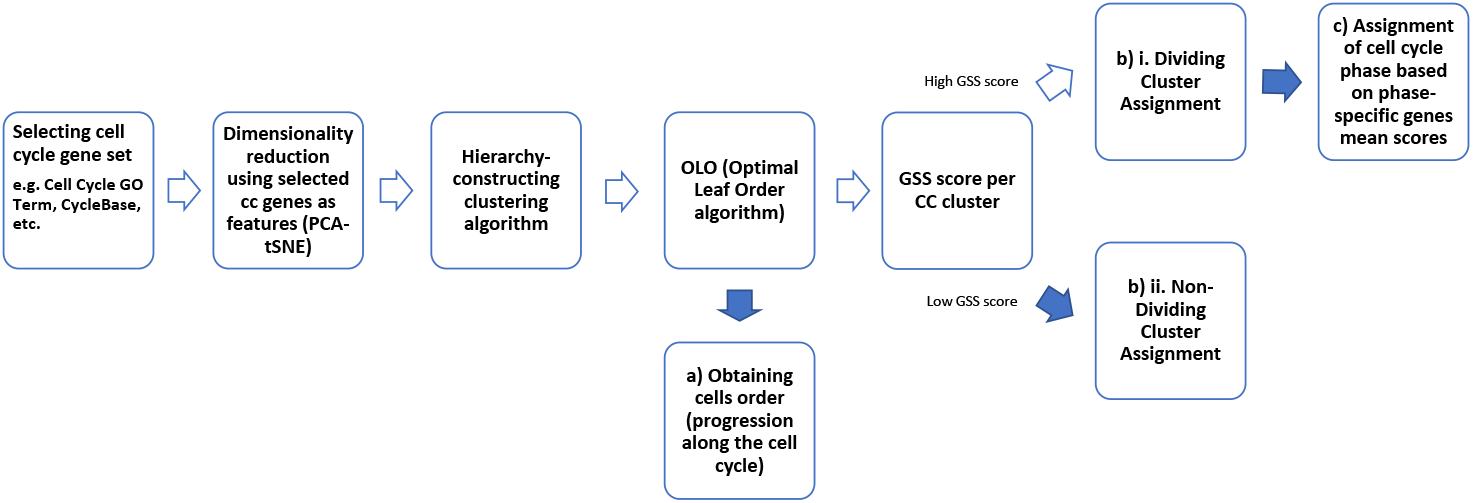
Process Flow Diagram of SC1CC. This simplified process flow shows a typical sequence of steps to analyze scRNA-Seq data with SC1CC; three outcomes can be obtained, a) the order of cells according to their progression along the cell cycle phases as determined by SC1CC, b) a designation of dividing vs. non dividing cell clusters, and c) the cell cycle phase annotation for the identified cell cycle clusters.

